# Pretty Good Yields allow the spatial management of multiple objectives in agricultural landscapes

**DOI:** 10.64898/2026.07.06.736684

**Authors:** Madeleine Kubasch, Manon Costa, Nicolas Loeuille

## Abstract

In order to feed a growing global population without silencing nature, conceiving agricultural management strategies reconciling yield and conservation goals is key. Using numerical simulations of a metacommunity model, we explore the possibilities for compromise offered by spatial management strategies of farmed areas. Each strategy is characterized by its farming intensity, the proportion of farmed lands and their spatial aggregation. We show that achieving equitable yield-biodiversity compromise is difficult. While conciliatory strategies offering top yield and biodiversity are typically not possible, accepting slightly lower yields (ie, “Pretty Good Yield strategies”) allows to recover substantial biodiversity. Such reconciliation possibilities are limited for species with small dispersal. Yield increases mainly through farmland expansion, whereas farming intensity strongly influences biodiversity, increasing it at low intensity before decreasing with further intensification. Finally, we demonstrate that reconciliation is easier if agricultural production relies on biodiversity through ecosystem services.

## 1. Introduction

Agriculture is at the heart of food security. With total global food demand being expected to rise by 35 to 56% between 2010 and 2050 (van Dijk et al. 2021), maintaining or possibly increasing farming yield appears essential. This however comes at the risk of biodiversity loss, as agricultural intensification is one of the main drivers of the current biodiversity crisis (Habel et al. 2019, Rigal et al. 2023). Indeed, farmland expansion leads to habitat loss and fragmentation, whereas conventional intensification relying on input of agrochemicals (pesticides, fertilizers…) qualitatively degrades habitat quality for farmed lands and surrounding habitats.

Despite this apparent opposition, conservation and yield goals are intimately intertwined, as agricultural production relies on various ecosystem services, such as pollination, soil maintenance and fertility, nutrient regulation or enemy control. The delivery of multiple ecosystem functions intrinsically relies on biodiversity, as different species are typically required to sustain separate functions under contrasted environmental conditions (Isbell et al. 2011). As a consequence, the negative feedback of conventional intensification on biodiversity ultimately leads to intensification traps, where the negative impact of biodiversity loss on yield outweighs the immediate yield of intensive farming (Bommarco et al. 2013, Burian et al 2024). This challenges the concept of conventional intensification, which attempts to replace biodiversity services by energy and agrochemical inputs (Tilman et al. 2001). Thus, achieving a sustainable reconciliation of yield and conservation goals appears to be crucial.

Agricultural activities significantly rearrange the spatial configuration of natural habitats, both through farmland expansion and intensification. Spatial management conendrum is exemplified by the land sharing-sparing debate (Green et al. 2005) that explicitly links production and conservation goals. It contrasts two management strategies for farmed landscapes in order to achieve a fixed yield goal. Land sparing aims at minimizing the farmed area through conventional intensification, allowing to keep intact as much natural habitat as possible, whereas land sharing favors agroecological or organic practices that limit the negative impacts of conventional intensification, but ultimately needs a larger surface to get the fixed yield. As a consequence, land sparing leads to farmed landscapes which rely on a relatively small, spatially aggregated, intensively farmed area; whereas land sharing consists in mosaics of natural and agroecologically farmed patches.

Empirical evidence indicates that the impact of either strategy on conservation goals depends on several factors such as species’ mobility and habitat preferences (Augustiny et al. 2025). For instance, land sparing may be more favorable for native species that strongly depend on specific habitats and for species with low dispersal capacities, whereas land sharing or hybrid strategies tend to be beneficial for generalist or mobile species. Land sharing appears to be better at preserving ecosystem services such as pollination and pest control, which is also supported by recent modeling studies (Grass et al. 2019, Oliveira-Xavier et al. 2025).

Nevertheless, the land sharing-sparing framework has also been met with criticism (Fischer at al. 2014). Importantly, focusing solely on land sharing or land sparing extremes is restrictive, as many hybrid strategies are conceivable and appear to be well suited to reach conservation goals. In addition, the land sharing-sparing debate insists on maintaining a given yield production. However, accepting some yield loss may allow for substantial biodiversity gains. For example, in fisheries, moving from maximum yield approaches to *pretty good yield* strategies (at least 80% of maximum yield) can typically allow large improvements in fished populations (Hilborn 2010). Such opportunities for compromise are unfortunately missed by the land sharing-sparing debate.

In the present work, we aim at exploring the biodiversity-yield trade-off by considering a large set of spatial agricultural management strategies, including but not limited to land sharing-sparing strategies, thus going beyond this framework. More precisely, each management strategy is characterized by its spatial heterogeneity through three parameters: the proportion of farmed area, its spatial aggregation and farming intensity. In order to assess the performance of these management strategies, we consider a metacommunity model (Leibold et al 2004). A metacommunity consists in a pool of interacting species whose habitat is composed of several localities or patches, connected by dispersal. As a consequence, emergent biodiversity depends on two forces (Mouquet and Loreau 2003, Leibold et al 2004) : (i) *species sorting* with species concentrating in patches where they are best adapted:; (ii) *mass effects* where dispersal from suitable patches may allow species to be present even in less favorable habitats.

Building on this theoretical framework, we investigate possibilities for reconciling yield and conservation goals. Reconciliation is expected to be difficult when loss of ecosystem services may be mitigated by intensive farming, as yield and biodiversity are likely optimized by very different strategies. Conversely, it should be easier when ecosystem services are irreplaceable, as maintaining biodiversity then is key to achieving high yields. We identify strategies that allow to reach such conciliation, when possible. We expect that it is achieved either by land sharing or land sharing-sparing mixtures, given the previous evidence on the good performance of hybrid strategies, and the importance of land sharing for ecosystem services maintained through mass effects. Finally, since dispersal is crucial in metacommunity models, these conclusions are likely to depend on species’ mobility. In particular, lower dispersal capacity should make species more vulnerable to habitat loss and/or degradation, further limiting spill-over effects and emergent ecosystem services.

## 2. Methods

### 2.1 Presentation of the metacommunity model Landscape construction

We consider a family of spatial management strategies of agricultural landscapes which is parametrized by three parameters: proportion of farmed surface (*p*_*A*_), spatial aggregation (*H*) and farming intensity (*I*). In particular, the obtained management strategies include land sharing and land sparing strategies, but also cover a broader range of strategies (Figure 1a).

**Figure 1.**
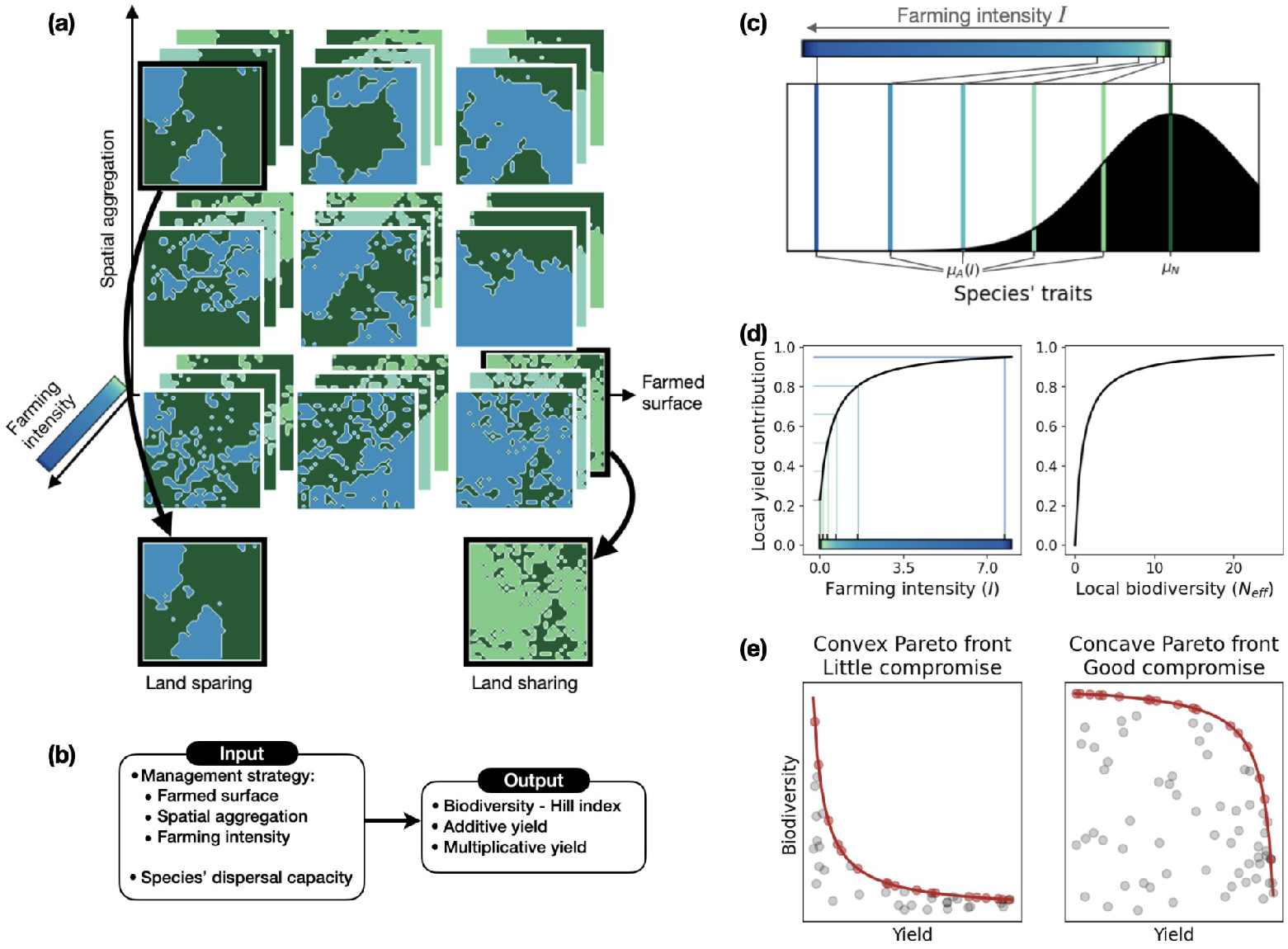
(a) Schematic representation of the explored space of agricultural landscapes, highlighting the three spatial management parameters under study. (b) Schematic representation of the model, indicating input parametersvaried throughout the simulation study, and model outputs. (c) Species trait distribution (black shadow) centered on traits μ_N_ that are adapted to natural habitats. Trait optima in agricultural patches are increasingly different when farming intensity I becomes high (see the differentcolored lines). (d) Respective contributions of farming intensity and local biodiversity to local yield. On the left panel, colored lines indicate the six levels of farming intensity which are subsequently used in simulations. (e) Schematic representation of the biodiversity-yield trade-off that contrast in terms of conciliation possibility. Each data point represents the average biodiversity and yield achieved by a management strategy. The Pareto front is shown in red.

The proportion of farmed surface varies on a regular grid of five values between 10% and 90%, to which we add the case of a totally natural landscape (*p*_*A*_ *= 1, I = 0*), and a totally farmed one (*p*_*A*_ *= 1, I > 0*). Spatial aggregation varies on a regular grid of five values between 0.01 (low aggregation) and 0.99 (high aggregation). Finally, we consider six farming intensity levels, from zero intensity (hunter-gatherer) to intensive farming. Farming intensity levels are fixed in such a way that its local yield contributions (see forthcoming Section 2.2 and Supplementary Information A.4, Equation S7) form a regular grid, from minimal yield (zero intensity) to 95% of maximum yield (Fig. 1d).

At a fixed proportion of farmed surface and spatial aggregation, a landscape realization is generated by a stochastic landscape model of tunable spatial aggregation (Supplementary Information A.1). Spatial aggregation is defined by the Hurst index *H ∈ (0,1)* which is such that spatial aggregation is high at low values of *H*, and vice-versa. The landscape model relies on a fractional Brownian sheet of Hurst index *H* (Etherington et al. 2015), and we refer to Supplementary Information A.1 for detail on its mathematical definition (Equation S1). For each combination of parameters *(p*_*A*_, *H)*, we generate 100 stochastic realizations of the agricultural landscape, which will be farmed at all intensity levels defined above (Fig. 1a).

From now on, we assume the agricultural landscape to cover the square *Ω = [0,1]*^*2*^, and designate by *A* and *N* the farmed and natural areas, respectively.

### Local habitat characteristics

In order to distinguish different ecological strategies, each species will be characterized by a trait which represents its adaptation to natural and farmed habitat. For each habitat type, there exists a trait value which is assumed to be optimal. More precisely, optimal trait values in natural and farmed habitat are respectively *μ*_*N*_ *= d/2* and *μ*_*A*_ *= -d/2*, where *d* designates the distance among optimal traits. A species is considered to be a generalist if their trait is close to zero, whereas it is a natural specialist (resp. agricultural specialist) if their trait is close to *μ*_*N*_ (resp. *μ*_*A*_). We expect the distance between optimal traits to grow as farming intensity increases, as habitats become increasingly different. For parsimony, we define the distance between optimal traits as a type II function of farming intensity. Detail is provided by Equation S2 in Supplementary Information A.2.

### Species trait distribution

We consider natural habitat as the starting point, the question being how this pool of species may maintain when agricultural development is undertaken. We thus assume that species traits are independent and identically distributed, following a normal distribution which is centered on *μ*_*N*_ (Fig. 1c). Its variance is chosen to ensure that with probability 95%, at least 5% of space is occupied at equilibrium by a metapopulation in a homogeneous environment (Equations S4 and S6, Supplementary Information A.3). In practice, we consider 100 independent, identically distributed species pools of 30 species each.

### Local extinction rates

Local extinction rates increase when species trait and local habitat type do not match. For *X ∈ {A,N}*, the local extinction rate of a species of trait *μ* in farmed (*X = A*) or natural (*X = N*) habitat is given by

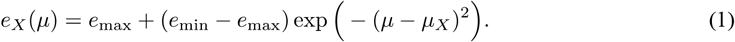

Consider species *j* of trait *μ*_*j*_. Its local extinction rate τ_j_(x) at any point *x ∈ Ω* is defined as

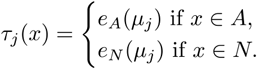

Minimal and maximal extinction rates *(e*_*min*_, *e*_*max*_) are chosen based on the proportion of occupied space at equilibrium by a metapopulation in a homogeneous environment. The maximal extinction rate is the lowest rate for which such a metapopulation goes extinct, whereas the minimal extinction rate ensures that it occupies 99% of space at equilibrium (Equations S4 and S5, Supplementary Information A.3).

### Colonization dynamics

A colonization attempt occurs whenever a species disperses from some point *x ∈ Ω* at which it is present to another point *y ∈ Ω*. We assume that colonization rates decrease with distance. For *x = (x*_*1*_ , *x*_*2*_*) ∈ Ω* and *y = (y*_*1*_ , *y*_*2*_*) ∈ Ω*, the rate at which species *j* disperses from *x* to *y* is thus defined by the following exponential dispersal kernel:

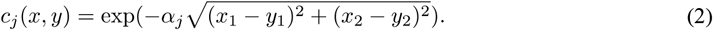

This dispersal kernel is parametrized by the average dispersal distance α_j_ ^-1^. Here, all species are assumed to have equal dispersal capacity, hence *α*_*j*_ *= α*. In order to explore the impact of the species’ dispersal capacity on the biodiversity-yield trade-off, we consider three levels of dispersal. Our baseline scenario of large dispersal corresponds to an average dispersal distance of 1.3 km in a landscape of 50 × 50 kilometers, yielding *α = 37*.*5*. This scenario is based on estimated dispersal distances for butterflies taking into account long-distance dispersal events (Stevens et al. 2010). Intermediate and small dispersal scenarios are obtained by dividing the average dispersal distance by a factor 1.75 or 2.5, respectively.

Let us assume that species *j* (with trait *μ*_*j*_) reaches some point *y ∈ Ω* by dispersal. If no species is present at *y*, then species *j* is able to establish itself at *y* and the colonization attempt is successful. However, if some species *i* (with trait *μ*_*i*_) is already present at *y*, the outcome depends on the competitive hierarchy between species. We assume local competitive exclusion, so that species *j* replaces species *i* at *y* if it is better adapted to the local habitat, *i*.*e*. if trait *μ*_*j*_ is closer to the habitat optimum than the trait *μ*_*i*_. Otherwise, species *i* remains present at *y*.

This competitive hierarchy between species *i* and *j* is summarized by a function *ϕ*_*ij*_, which equals one at any point *x* at which species *i* may replace species *j*, and zero elsewhere:

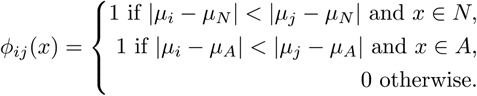

### Master equation of metacommunity dynamics

The ecological dynamics described above give rise to the following master equation, which arises from stochastic colonization-extinction dynamics by considering a large number of patches that are uniformly distributed throughout the landscape (Costa et al. 2026). We consider a set of *S* species, and aim to characterize the probability *u*_*j*_*(t,x)* of observing species *j* at location *x* at time *t*. The species are in competition for habitat space, and local competitive exclusion implies that

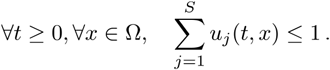

The metacommunity model is given by the following system of integro-differential equations. For any *j∈ {1,…, S}, t ≥ 0* and *x ∈ Ω*,

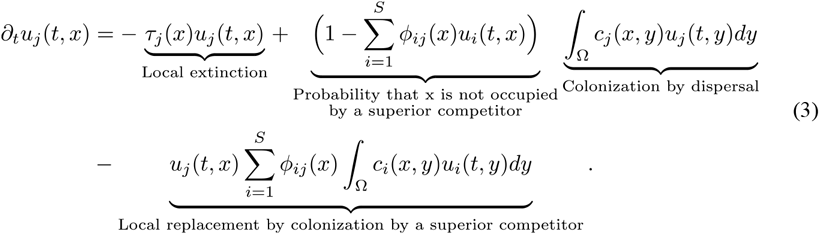

These equations are interpreted as follows. If species *j* is present at location *x*, it can become locally extinct at rate *τ*_*j*_*(x)*, depending on its adaptation to local conditions (match between the species trait and the local habitat). In addition, if species *j* is currently absent from *x*, it may attempt colonization from another location *y* which it occupies at rate *c*_*j*_*(x,y)*, provided that *x* is not currently occupied by a locally superior competitor. Finally, species *j* may be replaced by a locally superior competitor *i* at *x* through colonization from any point *y* at which species *i* is currently present. This occurs at rate *c*_*i*_*(x,y)*, which depends on the distance separating *x* from *y*.

### 2.2 Simulation study and analysis of model outputs

#### Simulation scenarios

For each each farming strategy *(p*_*A*_, *H, I)* and average dispersal distance *α*, we perform 100 simulations where the k-th simulation combines the k-th landscape realization with the k-th species pool.

The initial condition of the metacommunity model (3) is fixed to uniform species occupancies, equal to *1/(S+1)*. We let the simulation run for 5000 time units. In particular, we do not necessarily wait for the ecological dynamics to reach their equilibrium. Indeed, long transients may occur for instance if two species have very similar traits, in which case coexistence may be observed over very long time spans even though eventually only one species persists. Notably, such dynamics are relevant in our setting, as agricultural practices are conceived on relatively short timescales (Hastings 2004), compared to their ultimate consequences for biodiversity (eg, extinction debts, Tilman et al 1994).

The study is repeated for three different dispersal scenarios (Section 2.1), keeping all other parameters fixed. As we consider 156 farming strategies, this amounts to 46800 simulations. Figure 1b summarizes the relevant model inputs and outputs.

### Definition of model outputs

For food production, the key model output is farming yield. Here, we first define a local yield *ρ(t,x) ∈ [0,1]*, which depends both on farming intensity and on local biodiversity through ecosystem services (Fig 1d).

More precisely, the intensity-yield contribution is a type II function of farming intensity (Supplementary Information, Equation S7), whereas the biodiversity-yield contribution is a type II function of local effective species number (Supplementary Information, Equations S8 and S9). This allows to take into account saturating effects of intensification and biodiversity-ecosystem functioning (Loreau et al. 2001, Burian et al. 2024). We refer to Supplementary Information A.4 for detail.

We consider two different formulas for local yield, which embody different conceptions of farming intensification. Additive yield equals the sum of intensity and biodiversity contributions (Supplementary Information, Equation S10). This corresponds to a traditional point of view, where conventional intensification can replace ecosystem services. Alternatively, multiplicative yield is defined as the product of intensity and biodiversity contributions (Supplementary Information, Equation S11), which implies that ecosystem services cannot be replaced by conventional intensification. Total yield is then obtained by integrating local yield over the whole agricultural area:

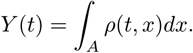

We measure biodiversity at metacommunity scale, as in *γ*-diversity. The indicator should further be sensitive to both species’ rareness, and to the probability of patches being unoccupied. In particular, classical biodiversity indicators like species richness and Shannon diversity do not take this information into account (Supplementary Information A.5, Equations S12 and S12). We therefore use a biodiversity indicator adapted from classical Hill indices (Hill 1973, Jost 2006; Supplementary Information, Equations S14 and S15). This index depends on the absolute abundance of species *j* averaged over the total landscape at time *t*, and is defined as follows:

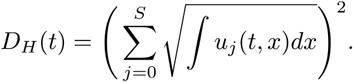

We refer to Supplementary Information A.5 for more detail on the properties of this biodiversity indicator.

### Conciliation analysis

For the rest of our analysis, we consider average values of yield and biodiversity obtained by averaging over the 100 simulations for each strategy and dispersal scenario. Importantly, our main focus is the analysis of qualitative characteristics of possible biodiversity-yield trade-off, depending on the dispersal level and biodiversity-yield relationship. As a consequence, at a given dispersal level, average yield and biodiversity values are normalized by the average values achieved in a homogeneous natural landscape.

In order to identify yield-biodiversity compromise opportunities, we consider an approach inspired by the Pretty Good Yield concept of Hilborn (2010). The idea is to allow some yield loss in order to recover some biodiversity: the higher the biodiversity gain, the easier reconciliation will be. Here, we consider strategies to be of Pretty Good Yield if their yield equals at least 75% of the highest observed yield. Reciprocally, we also define Pretty Good Biodiversity strategies, which achieve at least 75% of the highest observed biodiversity. In particular, strategies which are both of Pretty Good Yield and Pretty Good Biodiversity are considered as satisfactory compromises.

The biodiversity-yield trade-off is further investigated by considering the Pareto front of efficient strategies (Fig. 1e). A management strategy is Pareto-efficient if no other strategy achieves both better yield and better biodiversity. If the Pareto front is concave, conciliation is easier as accepting small yield losses leads to substantial biodiversity gains, and vice-versa. Conversely, conciliation is difficult for convex Pareto fronts, as substantial biodiversity gains are necessarily linked to substantial yield losses.

## 3. Results

### 3.1 Achieving a fair yield-biodiversity compromise is difficult, and strongly depends on species dispersal

Additive yield and biodiversity are optimized by very different strategies (Fig. 2). While additive yield is maximized by intensively farming the whole landscape (Fig 2g), biodiversity peaks when some natural habitat is preserved and farming intensity is low. More precisely, at high and medium dispersal, biodiversity is best for strategies that are close to land sharing (50% of farmed area at low intensity, low spatial aggregation; Fig. 2d), whereas it requires small, low intensity farmed lands which are highly aggregated at small dispersal (Fig. 2e). As a consequence, conciliation of conservation and production goals is difficult and trade-offs need to be considered.

**Figure 2.**
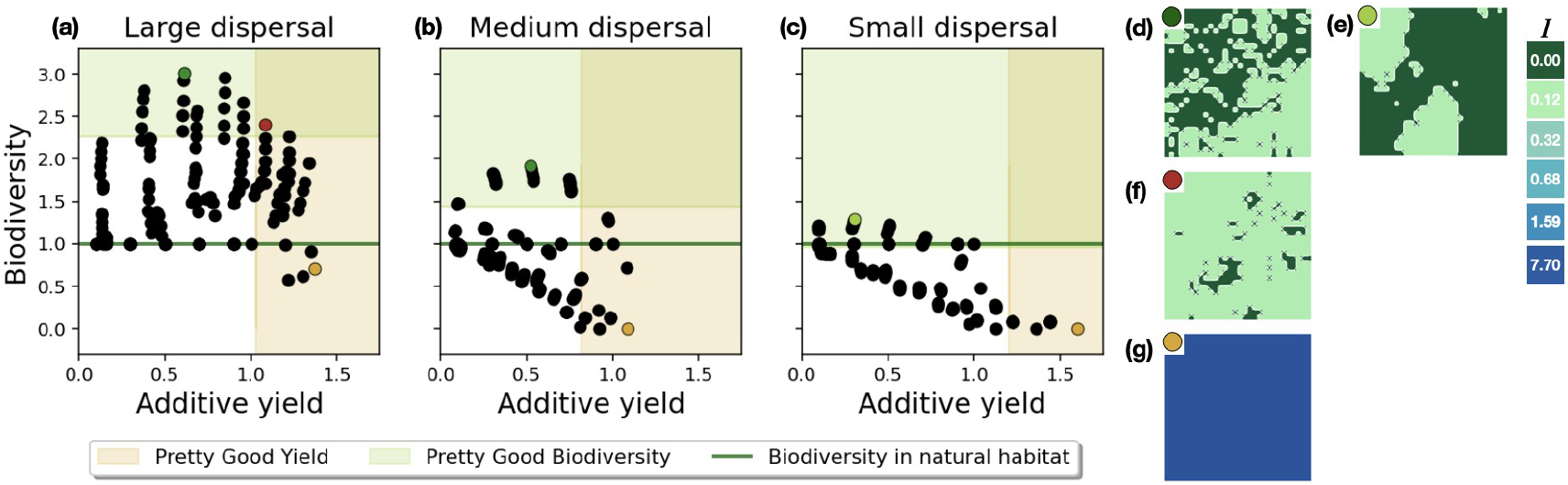
(a-c) Scatter plot of average values of additive yield and biodiversity obtained by each management strategy. Orange and green shaded areas correspond respectively to areas of pretty good yield (yield superior to 75% of maximal observed yield within the panel) and pretty good biodiversity (biodiversity superior to 75% of maximal observed biodiversity within the panel). The dark green line indicates the biodiversity level achieved in a homogeneous natural environment, which is normalized at one. (d-g) Landscapes maximizing biodiversity mix habitat types with low agricultural intensity and an aggregation that depends on species dispersal (d,e) or additive yield, here requiring a total, intense exploitation (g), and a possible conciliation at large dispersal that mixes natural habitats in a matrix dominated by low intensity farming (f).

Considering Pretty Good Yield (PGY) and Pretty Good Biodiversity (PGB) strategies provides significant leverage to reach such conciliation (Fig. 2a-2c). Conciliation strongly depends on species dispersal. At intermediate or high dispersal, the PGY and PGB strategies clearly intersect (Fig. 2a), and a compromise is achieved by management strategies devoting most of the landscape to low intensity unaggregated farming (Fig. 2b, 2f). At low dispersal, strong trade-offs occur between the two objectives. Most management strategies lead to significant biodiversity losses compared to a homogeneous natural habitat (Fig. 2c). PGY and PGB are never met simultaneously, which implies that either yield or conservation goals need to be prioritized as no conciliation is achievable.

### 3.2 Efficient strategies are located along a farmland expansion and intensification gradient

Additive yield and biodiversity mostly depend on contrasting parameters. Indeed, additive yield is increased mainly by expanding cultivated surfaces (horizontal color organization on Fig. 3a, 3d and 3g), whereas biodiversity is mainly harmed by enhancing intensity (vertical color organization on Fig. 3b, 3e, and 3h).When moving along the Pareto front from optimal biodiversity to optimal additive yield, both farming intensity and proportion of farmed area increase. The underlying management strategies are thus gradually intensified. Importantly, these conclusions still hold when taking into account the dispersal of these model outputs around their mean (Supplementary Information B.1, Fig. S2).

**Figure 3.**
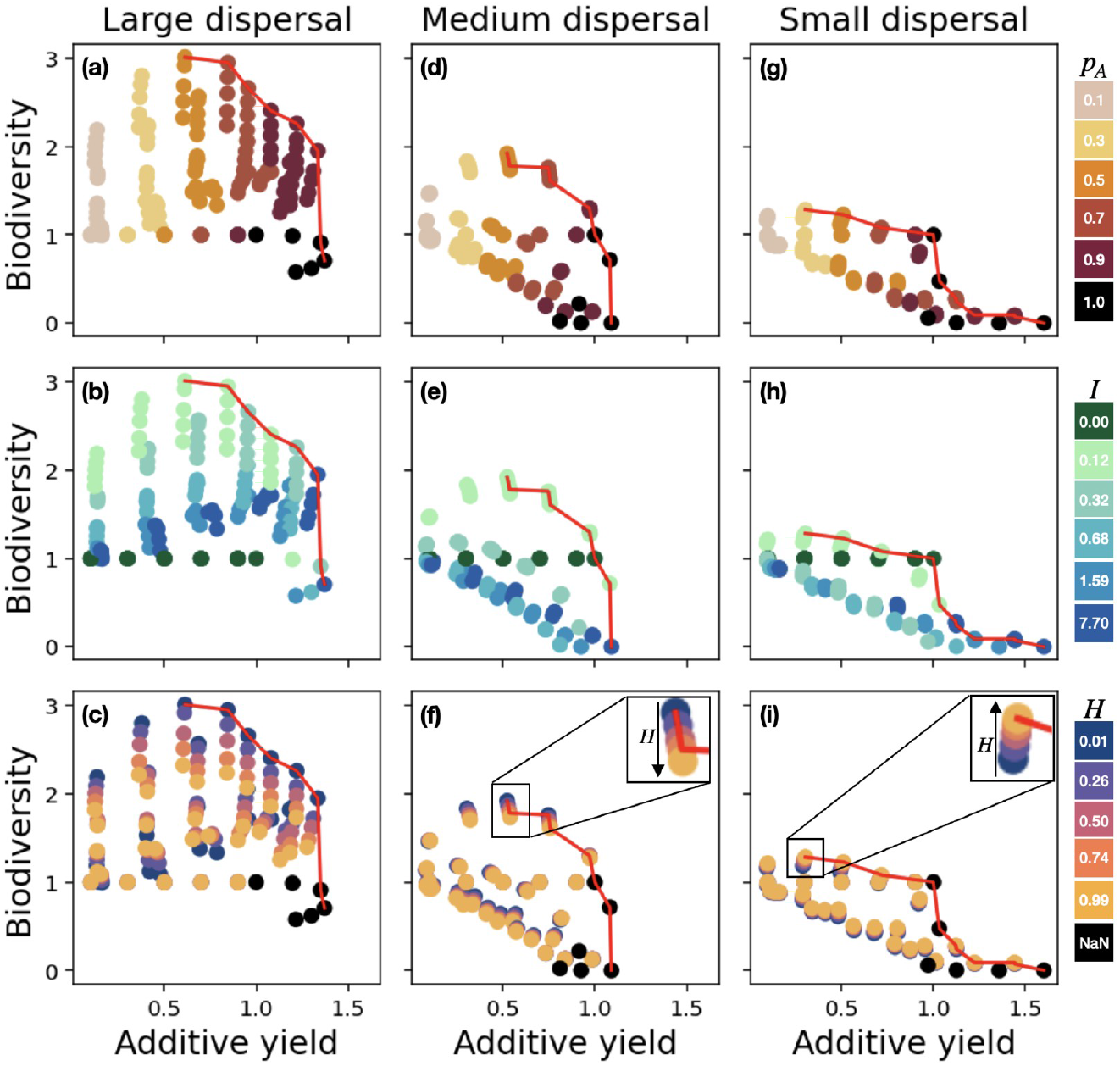
Average values of additive yield and biodiversity achieved by each management strategy, at large (a-c), medium (d-f) and small dispersal (g-i). Each management strategy is characterized by proportion of farmed area (a, d, g), farming intensity (b, e, h) and spatial aggregation (c, f, i). For homogeneous landscapes (p_A_ = 1), spatial aggregation is not defined (H = NaN). The Pareto front is shown in red.

While aggregation plays a lesser role (less straightforward color organization on fig 3c, f, I), it can offer substantial gains in biodiversity and facilitate conciliation. For a fixed surface/intensity scenario, at high and medium dispersal, decreasing spatial aggregation enhances biodiversity while keeping yield fixed (inset on Fig. 3f). At small dispersal, the same outcome is achieved by increasing spatial aggregation (inset on Fig. 3i). Notably, at high and medium dispersal, it is possible to increase both yield and biodiversity of some PGB strategies by simultaneously decreasing spatial aggregation and increasing farmed surface or farming intensity. These results not only stress the importance of spatial configuration, but also that conservation benefits for small and large dispersal species depend on alternative spatial strategies.

### 3.3 Reconciliation is easier when yield and biodiversity are strongly connected

When recognizing that ecosystem services and conventional intensification are not clearly substitutable, but in fact interact, conciliation is somewhat facilitated. Indeed, when comparing multiplicative yields to previous additive yields, we note that the Pareto front is moved right, and that more strategies fall in the conciliation part. While agricultural production still spreads over the whole landscape, low farming intensity is required (Fig. 4f, see also Fig. S3 for high and medium dispersal). Preserving biodiversity is then more beneficial and multiplicative yield and biodiversity break down simultaneously at high farming intensity (Supplementary Information B.2, Fig. S4). Accounting for the dispersal of outputs around their means does not alter these qualitative conclusions (Supplementary Information B.2, Fig. S5). Notably, conciliation always is achievable for multiplicative yield, even at small dispersal. Indeed, the Pareto front is concave: contrary to additive yield (Fig. 4a), accepting some multiplicative yield loss leads to significant biodiversity gains (Fig 4b). As a consequence, some strategies satisfy both PGY and PGB (Fig 4b), including the totally natural landscape used for production (hunter-gatherer strategy, Fig. 4d). Equitable trade-offs are thus feasible, though conciliation still concerns only few spatial configurations.

**Figure 4.**
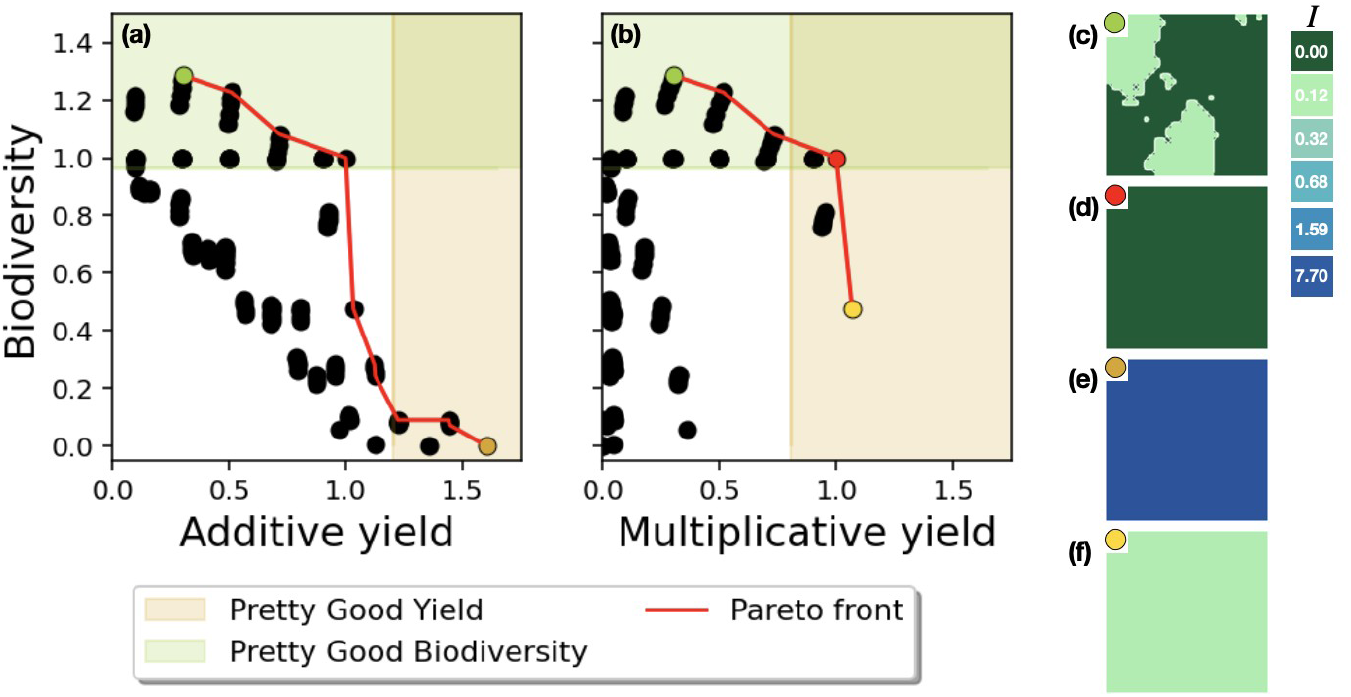
(a-b) Scatter plot of average values of additive (a) or multiplicative (b) yield and biodiversity obtained by each management strategy, at small dispersal. Orange and green shaded areas correspond respectively to areas of pretty good yield (yield superior to 75% of maximal observed yield) and pretty good biodiversity (biodiversity superior to 75% of maximal observed biodiversity). (c-f) Examples of landscapes maximizing biodiversity (c), additive yield (e), multiplicative yield (f), and a compromise strategy for multiplicative yield (d).

## 4. Discussion

Our results demonstrate that conciliation of production and conservation goals in agricultural landscapes is difficult, especially when ecosystem services may be replaced with conventional intensification (additive yields). Biodiversity is strongly impaired by the increase of farming intensity, whereas farmland expansion benefits yield without necessarily affecting biodiversity. Our trade-off analysis however highlights that accepting some yield and biodiversity loss allows allows conciliation, except for low dispersal species. When ecosystem services cannot be replaced by farming practices (multiplicative yield), conciliation is slightly easier, and possible even at low dispersal.

Going beyond the land sharing-land sparing debate proved crucial to identify relevant landscape management strategies for biodiversity-yield compromise. Indeed, most of the Pareto-efficient strategies correspond to intermediate management strategies (Fig. 3). While this has previously been argued in the literature (Grass et al. 2019; Oliveira-Xavier et al, 2025; Augustiny et al. 2025), to our knowledge, our study is the first exhaustive exploration of management strategies over all landscape parameters which intervene in the classical land sharing-sparing debate (agricultural expansion, intensification and spatial aggregation). We also stress that no single management strategy is optimal under all considered scenarios, neither for conservation, nor for production or conciliation. Indeed, optimal strategies depend on dispersal and/or the reliance of yield on ecosystem services. This is consistent with empirical studies favoring either land sharing, land sparing or mixed strategies depending on the system at hand (Augustiny et al. 2025). While land sparing never is optimal in our setting, land sharing or mixed strategies are crucial to sustain biodiversity and reach conciliation.

While the traditional land sharing-land sparing debate relies on fixed yields, our results highlight that allowing some yield loss can substantially broaden the set of conciliatory strategies. This part of the analysis is inspired by works on Pretty Good Yields in fisheries (Hilborn 2010). The idea of reducing agricultural production may appear in stark contrast to the expected rise in world-wide food demand. However, even in this context, a focus on yield is not optimal as several other aspects such as food security or sovereignty are also crucial (Fischer et al 2014). About one third of the food produced for human consumption is lost or wasted (FAO, 2011). Similarly, part of our food production actually exceeds what is strictly speaking required for human nutrition, for instance cocoa, coffee or animal feed (Fischer et al. 2014). Changes in food consumption, such as reduced-meat diets, could thus further lower the urge to increase agricultural yield itself.

Our findings further emphasize that strong dependency of agricultural production en ecosystem services improves conciliation. In our setting, additive yield indeed corresponds to a conventional viewpoint on agricultural intensification: farming practices and agrochemical inputs replace ecosystem services. However, while some ecosystem services such as nutrient supply have *de facto* been replaced by fertilizers in conventional farming systems (Fontaine et al. 2024), other ecosystem services like pollination are hard to substitute (Broussard et al. 2023). When conventional intensification successfully increases yields, its negative impact on biodiversity is well-documented (Tscharntke et al. 2005, Habel et al. 2019, Rigal et al. 2023), and our results emphasize that reconciling yield and biodiversity indeed is difficult in this context. Ultimately, this can create a feedback loop where increasing farming intensity impairs irreplaceable ecosystem services, requiring further intensification (“intensification traps”, Burian et al. 2023). A shift in paradigm has thus been proposed, harvesting the benefits of ecosystem services by appropriate management of the underlying agricultural landscapes (Bommarco et al. 2013, Tscharntke et al 2021). Our study accounts for such ecological intensification through multiplicative yields. This pathway is pertinent for biodiversity-yield conciliation, as it consistently enables equitable trade-offs (Figure 4). Nevertheless, biodiversity and yield are still optimized by very different strategies: in order to protect biodiversity, landscape management thus needs to actively account for conservation goals. This is reminiscent of previous findings that maximizing ecosystem services may not necessarily lead to actually preserving biodiversity (Macfayden et al., 2012).

While biodiversity is affected most strongly by farming intensity, the qualitative impact of farmland expansion on biodiversity depends on the intensity level (Figure 3). At high intensity, increased proportion of space allocated to agriculture directly translates into habitat loss for most species, which in turn leads to biodiversity loss. This result is expected, based on previous investigations (Fahrig, 2003, Tscharntke et al. 2021, Bishop et al 2025). At intermediate farming intensity, the agricultural portion no longer makes for habitat loss, but for a somewhat hospitable matrix. This is known to increase biodiversity, both through theoretical model predictions and empirical studies (Murphy and Lovell-Dust, 2004; Driscoll et al. 2013; Suarez-Castro et al. 2026). In particular, a more hospitable matrix promotes effective connectivity between patches, as mortality in the matrix decreases. This likely is relevant in our model, as local extinction rates of species that are well adapted to natural habitat increase with farming intensity in agricultural habitat. Finally, at low farming intensity, farmed lands are actually viable habitat for most species, but dominant species differ between both habitat types. As a consequence, introducing some low-intensity farmed habitat actually creates a new niche, which allows for species sorting. This is beneficial for biodiversity in all scenarios (Figure 3). Such benefits of agroecosystems for biodiversity are well documented and recognized for some traditional farming practices, *e*.*g*. High Nature Value farmland in Europe (Tscharntke et al. 2005, Lomba et al. 2020, Tscharntke et al. 2021). This positive effect peaks at intermediate proportions of farmed area. This non-linear relationship is consistent with the “habitat heterogeneity hypothesis” (Pianka 1966) as well as predictions from source-sink metacommunity models (Mouquet and Loreau 2003), and has previously been observed in past agroecosystems through empirical studies of pollen records (Gordon et al. 2026).

Effects of spatial aggregation on biodiversity are comparatively weaker than the impact of farmland intensification or expansion. Here, for a given surface devoted to agriculture, higher aggregation makes for lower fragmentation “per se” (Fahrig 2017). Our results are therefore in line with the fact that effects of habitat fragmentation *per se* may be smaller that those of habitat loss (Fahrig, 2003). Nevertheless, habitat fragmentation does influence biodiversity in our model. While the direction of its residual effect on biodiversity is much discussed in the literature (Fahrig 2017, Fletcher et al 2018a, Fahrig et al 2019, Valente et al 2023), here it is clearly positive for good dispersers and negative for poor dispersers. This may be due to good dispersers being less sensitive to environmental perturbations at relatively small spatial scales (Tscharntke et al. 2005) and recalls that negative impacts of fragmentation occur at the same spatial scale as species dispersal (Fletcher et al. 2018b).

Importantly, these results on aggregation highlight that optimal strategies for biodiversity conservation and conciliation differ depending on species dispersal capacity. Several groups (birds, insects, amphibians…) are often used to discuss conservation measures in agricultural landscapes (Augustiny et al. 2025). As they have different dispersal abilities, a given strategy may impact one group more strongly than another. In particular, if an optimal solution is found based on the study of only one group, it may not be optimal for others, leading to possible trade-offs among conservation goals. Similarly, ecosystem services may rely on groups that have different dispersal ranges, including soil engineers, pollinating insects and pest regulating birds or mammals (Bommarco et al. 2013). This again makes emerge trade-offs among different ecosystem services. As a consequence, the conception of agricultural landscapes which sustain species’ that have different dispersal capacities is a key possible future perspective of our work.

Conceiving management strategies of agricultural landscapes that reconcile production and conservation goals is key to succeed in feeding humanity without silencing nature. Our results contribute to the identification of possible leverages which allow to reach equitable biodiversity-yield trade-offs. We emphasize that accepting some yield and biodiversity loss is necessary to reach a sustainable compromise, and that conciliation opportunities exist if yield strongly depends on ecosystem services, as in ecological intensification. We also highlight the importance of species dispersal range when managing agricultural landscapes. Our results thus participate in the on-going effort to transcend the classical land sharing-sparing debate (Fischer et al. 2014, Grass et al. 2019, Augustiny et al. 2025), and reveal crucial research questions on emerging multiple trade-offs between several conservation goals or ecosystem services.

## Acknowledgements

The simulations were performed on the SACADO MeSU platform at Sorbonne Université. This work has been supported by the Chair “Modélisation Mathématique et Biodiversité” of Veolia Environnement-Ecole Polytechnique-Museum national d’Histoire naturelle-Fondation X, and by ANR project HAPPY (ANR-23-CE40-0007).

## Statement of authorship

All authors conceived the model and contributed to the interpretation of results. Madeleine Kubasch performed the analysis and wrote the first draft. All authors made corrections and modifications, and approve the submitted version.

### Data accessibility

Scripts used to perform simulations, model output analysis and produce figures (main and supplementary) are available at https://gitlab.com/m-kubasch/metacommunity-simulations

## Bibliography

Aizen, N., Sáez, A., Morales, C.L. & Aizen, M.A. (2026). The conventional-to-organic yield gap diminishes with increasing crop pollinator dependence. Proc Biol Sci, 293, 20252553.

Augustiny, E., Frehner, A., Green, A., Mathys, A., Rosa, F., Pfister, S., et al. (2025). Empirical evidence supports neither land sparing nor land sharing as the main strategy to manage agriculture– biodiversity tradeoffs. PNAS Nexus, 4, pgaf251.

Barnsley, M.F., Devaney, R.L., Mandelbrot, B.B., Peitgen, H.-O., Saupe, D. & Voss, R.F. (1988). The Science of Fractal Images. Springer, New York, NY.

Bishop, G.A., Kleijn, D., Albrecht, M., Bartomeus, I., Isaacs, R., Kremen, C., et al. (2025). Critical habitat thresholds for effective pollinator conservation in agricultural landscapes. Science, 389, 1314–1319.

Bommarco, R., Kleijn, D. & Potts, S.G. (2013). Ecological intensification: harnessing ecosystem services for food security. Trends in Ecology & Evolution, 28, 230–238.

Broussard, M.A., Coates, M. & Martinsen, P. (2023). Artificial Pollination Technologies: A Review. Agronomy, 13, 1351.

Burian, A., Kremen, C., Wu, J.S.-T., Beckmann, M., Bulling, M., Garibaldi, L.A., et al. (2024). Biodiversity–production feedback effects lead to intensification traps in agricultural landscapes. Nat Ecol Evol, 8, 752–760.

Costa, M., Kubasch, M. & Loeuille, N. (2026) “Metacommunity persistence on spatially heterogeneous landscapes”, personal communication.

van Dijk, M., Morley, T., Rau, M.L. & Saghai, Y. (2021). A meta-analysis of projected global food demand and population at risk of hunger for the period 2010–2050. Nat Food, 2, 494–501.

Driscoll, D.A., Banks, S.C., Barton, P.S., Lindenmayer, D.B. & Smith, A.L. (2013). Conceptual domain of the matrix in fragmented landscapes. Trends in Ecology & Evolution, 28, 605–613.

Etherington, T.R., Holland, E.P. & O’Sullivan, D. (2015). NLMpy: a python software package for the creation of neutral landscape models within a general numerical framework. Methods in Ecology and Evolution, 6, 164–168.

Fahrig, L. (2003). Effects of Habitat Fragmentation on Biodiversity. Annual Review of Ecology, Evolution, and Systematics, 34, 487–515.

Fahrig, L. (2017). Ecological Responses to Habitat Fragmentation Per Se. Annu. Rev. Ecol. Evol. Syst., 48, 1–23.

Fahrig, L., Arroyo-RodrÍguez, V., Bennett, J.R., Boucher-Lalonde, V., Cazetta, E., Currie, D.J., et al. (2019). Is habitat fragmentation bad for biodiversity? Biological Conservation, 230, 179–186.

FAO. (2011). Global food losses and food waste – Extent, causes and prevention. Presented at the International Congress Save Food! Food and Agriculture Organization of the United Nations, Rome.

Fischer, J., Abson, D.J., Butsic, V., Chappell, M.J., Ekroos, J., Hanspach, J., et al. (2014). Land Sparing Versus Land Sharing: Moving Forward. Conservation Letters, 7, 149–157.

Fletcher, R.J., Didham, R.K., Banks-Leite, C., Barlow, J., Ewers, R.M., Rosindell, J., et al. (2018a). Is habitat fragmentation good for biodiversity? Biological Conservation, 226, 9–15.

Fletcher, R.J., Reichert, B.E. & Holmes, K. (2018b). The negative effects of habitat fragmentation operate at the scale of dispersal. Ecology, 99, 2176–2186.

Fontaine, S., Abbadie, L., Aubert, M., Barot, S., Bloor, J.M.G., Derrien, D., et al. (2024). Plant–soil synchrony in nutrient cycles: Learning from ecosystems to design sustainable agrosystems. Global Change Biology, 30, e17034.

Gordon, J.D., Fagan, B., Finch, J., Gillson, L., Milner, N. & Thomas, C.D. (2026). Black Death Land Abandonment Drove European Diversity Losses. Ecology Letters, 29, e70325.

Grass, I., Loos, J., Baensch, S., Batáry, P., Librán-Embid, F., Ficiciyan, A., et al. (2019). Land-sharing/-sparing connectivity landscapes for ecosystem services and biodiversity conservation. People and Nature, 1, 262–272.

Green, R.E., Cornell, S.J., Scharlemann, J.P.W. & Balmford, A. (2005). Farming and the Fate of Wild Nature. Science, 307, 550–555.

Habel, J.C., Ulrich, W., Biburger, N., Seibold, S. & Schmitt, T. (2019). Agricultural intensification drives butterfly decline. Insect Conservation and Diversity, 12, 289–295.

Hastings, A. (2004). Transients: the key to long-term ecological understanding? Trends in Ecology & Evolution, 19, 39–45.

Hilborn, R. (2010). Pretty Good Yield and exploited fishes. Marine Policy, 34, 193–196.

Hill, M.O. (1973). Diversity and Evenness: A Unifying Notation and Its Consequences. Ecology, 54, 427–432.

Isbell, F., Calcagno, V., Hector, A., Connolly, J., Harpole, W.S., Reich, P.B., et al. (2011). High plant diversity is needed to maintain ecosystem services. Nature, 477, 199–202.

Jost, L. (2006). Entropy and diversity. Oikos, 113, 363–375.

Jost, L. (2007). Partitioning Diversity into Independent Alpha and Beta Components. Ecology, 88, 2427–2439.

Kisdi, É. (2002). Dispersal: Risk Spreading versus Local Adaptation. The American Naturalist, 159, 579–596.

Leibold, M.A., Holyoak, M., Mouquet, N., Amarasekare, P., Chase, J.M., Hoopes, M.F., et al. (2004). The metacommunity concept: a framework for multi-scale community ecology. Ecology Letters, 7, 601–613.

Lomba, A., Moreira, F., Klimek, S., Jongman, R.H., Sullivan, C., Moran, J., et al. (2020). Back to the future: rethinking socioecological systems underlying high nature value farmlands. Frontiers in Ecol & Environ, 18, 36–42.

Loreau, M., Naeem, S., Inchausti, P., Bengtsson, J., Grime, J.P., Hector, A., et al. (2001). Biodiversity and Ecosystem Functioning: Current Knowledge and Future Challenges. Science, 294, 804–808.

Macfadyen, S., Cunningham, S.A., Costamagna, A.C. & Schellhorn, N.A. (2012). Managing ecosystem services and biodiversity conservation in agricultural landscapes: are the solutions the same? Journal of Applied Ecology, 49, 690–694.

Mouquet, N. & Loreau, M. (2003). Community Patterns in Source-Sink Metacommunities. The American Naturalist, 162, 544–557.

Murphy, H.T. & Lovett-Doust, J. (2004). Context and connectivity in plant metapopulations and landscape mosaics: does the matrix matter? Oikos, 105, 3–14.

Oliveira-Xavier, A., Calmé, S. & Gravel, D. (2025). The land-blending strategy: Contribution of metapopulation theory to the land sparing-sharing debate. Land Use Policy, 155, 107577.

Pianka, E.R. (1966). Latitudinal Gradients in Species Diversity: A Review of Concepts. The American Naturalist, 100, 33–46.

Rigal, S., Dakos, V., Alonso, H., Auniņš, A., Benkő, Z., Brotons, L., et al. (2023). Farmland practices are driving bird population decline across Europe. Proceedings of the National Academy of Sciences, 120, e2216573120.

Siopa, C., Carvalheiro, L.G., Castro, H., Loureiro, J. & Castro, S. (2024). Animal-pollinated crops and cultivars—A quantitative assessment of pollinator dependence values and evaluation of methodological approaches. Journal of Applied Ecology, 61, 1279–1288.

Stevens, V.M., Turlure, C. & Baguette, M. (2010). A meta-analysis of dispersal in butterflies. Biological Reviews, 85, 625–642.

Suarez-Castro, A.F., Hajian-Forooshani, Z., Barbosa, M.P.B., Damasceno, G., Grenié, M., Ocampo-Peñuela, N., et al. (2026). Trait-explicit approaches cast new light on fragmentation’s effects on biodiversity. Trends in Ecology & Evolution, 41, 148–157.

Tilman, D., Fargione, J., Wolff, B., D’Antonio, C., Dobson, A., Howarth, R., et al. (2001). Forecasting Agriculturally Driven Global Environmental Change. Science, 292, 281–284.

Tilman, D., May, R.M., Lehman, C.L. & Nowak, M.A. (1994). Habitat destruction and the extinction debt. Nature, 65–66.

Tscharntke, T., Grass, I., Wanger, T.C., Westphal, C. & Batáry, P. (2021). Beyond organic farming – harnessing biodiversity-friendly landscapes. Trends in Ecology & Evolution, 36, 919–930.

Tscharntke, T., Klein, A.M., Kruess, A., Steffan-Dewenter, I. & Thies, C. (2005). Landscape perspectives on agricultural intensification and biodiversity – ecosystem service management. Ecology Letters, 8, 857–874.

Valente, J.J., Gannon, D.G., Hightower, J., Kim, H., Leimberger, K.G., Macedo, R., et al. (2023). Toward conciliation in the habitat fragmentation and biodiversity debate. Landsc Ecol, 38, 2717–2730.

