## Supplementary Information for "Pretty Good Yields allow the spatial management of multiple objectives in agricultural landscapes"

### A. Detail on the metacommunity model

#### A.1 Landscape model

In order to obtain landscapes of given proportion of farmed surface $p_A$ and spatial aggregation $H$, we proceed as follows.

We consider a fractional Brownian sheet $V_H$ of Hurst index $H \in (0,1)$ (Mandelbrot and Van Ness, 1968; Wu and Xiao, 2007), which is defined as a real-valued, centered Gaussian random field with the following covariance function. For any $x = (x_1, x_2) \in \mathbb{R}^2$ and $y = (y_1, y_2) \in \mathbb{R}^2$,

$$\mathbb{E}[V_H(x)V_H(y)] = \frac{1}{2}\Big((x_1^2 + x_2^2)^H + (y_1^2 + y_2^2)^H - \big((x_1 - y_1)^2 + (x_2 - y_2)^2\big)^H\Big). \qquad \text{(S1)}$$

In particular, for $H = 1/2$, we recover a Brownian field, whereas increments are negatively correlated while they are positively correlated for $H > 1/2$.

It remains to distinguish natural and farmed areas. For a given realization of $V_H$ on $[0,1]^2$, let $q_\alpha$ designate the empirical quantile of order α. The agricultural area is defined as

$$A = \{x \in [0,1]^2 : V_H(x) \geq q_{1-p_A}\},$$

whereas the remaining habitat is considered to be natural.

In practice, simulations of the fractional Brownian sheet are obtained with the NLMpy package, which implements the mid-point displacement algorithm (Etherington et al. 2015).

#### A.2 Distance between optimal traits

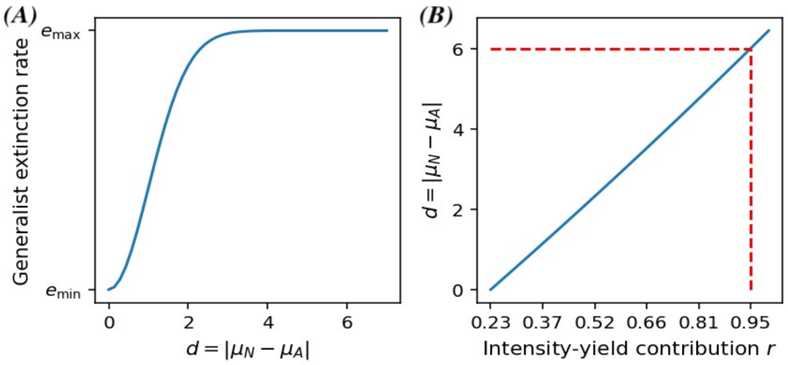


***Figure S1.*** *(A) Local extinction rate of the generalist species with trait* x = 0 *in both natural and farmed habitat, as a function of the distance* d *between optimal traits* $\mu_A$ *and* $\mu_N$ *in farmed and natural habitat. (B)* d *as a function of the intensity-yield contribution (Equation S3), for* $A_d = 11.35$ *and* $B_d = 1.76$.

We expect the distance between optimal traits to grow as farming intensity increases, as habitats become more and more different. For parsimony, we express the distance $d = |\mu_A - \mu_N|$ between optimal traits as the following type II function of farming intensity $I$:

$$d(I) = \frac{A_d I}{1 + B_d I} \tag{S2}$$

with $A_d, B_d > 0$ constants to be calibrated.

In order to fix $(A_d, B_d)$, we make the following assumptions. First, we consider that at high farming intensity, farmed and natural habitat are too different for generalist species to thrive. In other words, at high farming intensity, the local extinction rate of a species of trait $x = 0$ should be close to the maximal extinction rate $e_{max}$ (see forthcoming Section A3) in both natural and farmed habitats. As the highest farming intensity considered is approximately $I = 7.7$, we let $d(7.7) = 6$ (Figure S1, panel A).

Second, we expect that increasing farming intensity should have a similar impact on $d$ and on the intensity-yield contribution (Section A.4). Thus $d$ should be a roughly linear function of the local intensity-yield contribution $r \in (0.23, 1)$. Injecting forthcoming Equation (S5) into Equation (S1) provides the following expression of the distance between optimal habitats as a function of local intensity-yield contribution $r$:

$$d = d_y(r) = \frac{A_d(r - 0.23)}{1.87(1-r) + B_d(r - 0.23)}. \tag{S3}$$

As illustrated by panel B of Figure S1, letting $A_d = 11.35$ and $B_d = 1.76$ allows to satisfy these assumptions.

##### A.3 Local extinction rates and trait distribution

***Extremal extinction rates.*** In order to pick the minimal and maximal extinction rates, we consider a metapopulation in a homogeneous environment of dispersal kernel $c$ defined in Equation (2), and constant local extinction rate $\tau$. Analogously to Equation (3), the probability $w(t, x)$ that the species is present at location $x$ and time $t$ satisfies

$$\partial_t w(t,x) = -\tau w(t,x) + (1 - w(t,x)) \int_\Omega c(x,y) w(t,y) dy. \tag{S4}$$

The minimal and maximal extinction rates $(e_{min}, e_{max})$ are defined as the unique values of $\tau$ for which the metapopulation occupies respectively 99% or 1% of space at equilibrium. Their precise values are determined numerically, leading to

$$e_{\max} = 0.00354 \quad \text{and} \quad e_{\min} = 0.00003. \tag{S5}$$

***Trait distribution variance.*** We assume that the species traits are sampled from a normal distribution where $N(\mu_N, \sigma)$ is such that with probability 95%, a metapopulation in a homogeneous natural environment occupies at least 5% of space at equilibrium. Proceeding as described above, it follows that with probability 95%, the species extinction rates needs to be less than $e_Q = 0.00334$. Letting $q_\alpha$ designate the α-quantile of a centered reduced normal distribution, it follows from Equation (1) that

$$\sigma = \frac{\sqrt{2(\ln(e_{\max} - e_{\min}) - \ln(e_{\max} - e_Q))}}{q_{0.975}}. \tag{S6}$$

The trait distribution is thus properly characterized.

#### A.4 Local yield

Local yield *ρ(t, x)* results from a combination of intensity-yield and biodiversity-yield contributions. The intensity-yield contribution is defined as follows:

$$r(I) = r(0) + (1 - r(0))\frac{aI}{1 + aI}. \tag{S7}$$

Indeed, even without any intensification ($I = 0$), some yield is expected (hunter-gather scenario). In addition, we expect the intensity-yield contribution to saturate at high values of yield. Here, *r(0)* has been chosen in order to observe the same minimal intensity-yield contribution as in the meta-analysis conducted by Burian et al. (2024), thus $r(0) = 0.23$. Similarly, *a* equals the derivative at 0 of the intensity-yield contribution in Burian et al. (2024), leading to $a = 1.87$.

The biodiversity-yield contribution depends on the probability that *x* is occupied at time *t*, and the effective number $N_{eff}(t, x)$ of species at *x*. Let

$$u_0(t, x) = 1 - \sum_{j=1}^{S} u_j(t, x)$$

designate the probability of *x* not being occupied by any species at time *t*. Subsequently, given that *x* is occupied, the probability that species *j* is present at *x* is $v_j(t, x) = u_j(t, x)/(1 - u_0(t,x))$. The effective number of species (Jost et al 2007) hence equals

$$N_{\text{eff}}(t, x) = \exp\Big(-\sum_{j=1}^{S} v_j(t, x)\ln(v_j(t, x))\Big). \tag{S8}$$

Let us now turn to biodiversity-yield contribution. If no species is present at *x*, then the biodiversity-yield contribution is zero. Else, building on biodiversity-ecosystem functioning theory (Loreau et al. 2001), we expect it to be a type II function of local biodiversity, as measured by $N_{eff}$. At any time *t*, we thus define the biodiversity-yield contribution at *x* by

$$b(t, x) = (1 - u_0(t, x))\frac{N_{\text{eff}}(t, x)}{1 + N_{\text{eff}}(t, x)}. \tag{S9}$$

Finally, for additive yield, local yield is defined as

$$\rho(t, x) = (r(x) + b(t, x))/2 \tag{S10}$$

whereas multiplicative yield relies on

$$\rho(t, x) = r(x)b(t, x). \tag{S11}$$

Notice that in both cases, local yield takes values less than or equal to one.

#### A.5 Biodiversity indicators

In this section, we briefly discuss our choice of biodiversity indicator. As mentioned in the main text, our aim is to consider a regional biodiversity indicator which is sensitive to species' rareness, and to the risk that some spots may be unoccupied in our model with probability

$$u_0(t,x) = 1 - \sum_{i=1}^{S} u_i(t,x).$$

Usually, biodiversity indexes are defined as a function of the relative abundances of each species $i$ at $x$, which satisfy $\Sigma_{1 \leq i \leq S} v_i(x) = 1$ (Jost 2007, Roswell et al 2021). For instance, the regional $\gamma$-diversity based on Shannon diversity is defined by

$$D_\gamma = \exp\Big( - \sum_{i=1}^{S} \langle v_i \rangle \ln \langle v_i \rangle \Big) \tag{S12}$$

where

$$\langle v_i \rangle = \int_\Omega v_i(x) dx.$$

Let us consider an adaptation of $\gamma$-diversity to our model. We measure biodiversity at a fixed time $T$, and let

$$v_i(x) = \frac{u_i(T,x)}{1 - u_0(T,x)}$$

be the probability that at time $T$, species $i$ is present at $x$, given that $x$ indeed is occupied. We then may define $D_\gamma$ as in Equation (S3), and weight it by the average proportion of occupied space $1 - \langle u_0 \rangle$, leading to:

$$D_{\gamma,0} = (1 - \langle u_0 \rangle) D_\gamma. \tag{S13}$$

While this definition is parsimonious, it unfortunately is flawed, as illustrated by the following example. Assume that the agricultural habitat $A$ of total surface $p_A$ is occupied by exactly one species, which is absent from natural habitat, *i.e.* $u_1(T, x) = 1$ if $x \in A$ and $u_1(T, x) = 0$ otherwise. The latter instead is occupied uniformly by $S$ species such that for any $j \in \{2, ..., S + 1\}$, we have $u_j(T, x) = S^{-1}\varepsilon$ for a given $\varepsilon > 0$ if $x \in N$, and $u_j(T, x) = 0$ otherwise. A brief computation leads to

$$D_{\gamma,0}(\varepsilon) = (p_A + (1 - p_A)\varepsilon) \exp\left( - p_A \ln p_A - (1 - p_A) \ln\Big(\frac{1 - p_A}{S}\Big)\right),$$

where we emphasize the dependence in $\varepsilon$ as we will be interested in the behavior of this biodiversity index as $\varepsilon$ goes to zero. Indeed, in this case, we expect $D_{\gamma,0}$ to converge to its value if the natural habitat were totally unoccupied, which equals $p_A \exp(-p_A \ln p_A)$. However,

$$\lim_{\varepsilon \to 0} D_{\gamma,0}(\varepsilon) = p_A \exp\left( - p_A \ln p_A - (1 - p_A) \ln\Big(\frac{1 - p_A}{S}\Big)\right) > p_A \exp(-p_A \ln p_A).$$

This discrepancy is likely due to the fact that $D_{\gamma,0}$ separates the risk of spots being unoccupied from the information provided by the local relative abundances conditioned on occupation.

In order to correct for this flaw, one possibility is to consider a local biodiversity index, which subsequently is integrated over Ω in order to obtain a regional index. As an attempt to achieve this, we consider the local effective species number, conditioned on occupation:

$$N_{\text{eff}}(x) = \exp\Big( - \sum_{i=1}^{S} v_i(x) \ln v_i(x) \Big),$$

which subsequently is weighted by $u_0$ to obtain the following regional biodiversity index:

$$D_w = \int_\Omega (1 - u_0(T,x)) N_{\text{eff}}(x) dx.$$

However, this index again is not entirely satisfactory. Indeed, if we assume that only one species is present in the landscape such that $u_1(T, x) = 1$ for any $x$, we obtain as expected that $D_\omega = 1$. However, if we now assume that there are two species such that $u_1(T, x) = 1$ if $x \in A$ and $u_1(T, x) = 0$ otherwise, and $u_2(T, x) = 1$ if $x \in N$ and $u_1(T, x) = 0$ otherwise, we also have $D_\omega = 1$ despite the regional biodiversity being arguably higher in that scenario.

Finally, another possibility is to work directly with the species densities (or absolute abundances) $u_i$, instead of the relative abundances conditioned on occupation $v_i$. In order to define an appropriate biodiversity index based on species densities, we have considered the family of Hill indexes (Hill 1973, Jost 2006). These usually are defined for $l \in \mathbb{R}$ by

$$D_\ell = \Big( \sum_{i=1}^{S} \langle v_i \rangle^{1-\ell} \Big)^{1/\ell}, \tag{S14}$$

with the special case $l = 0$ given by

$$D_0 = \exp\Big( - \sum_{i=1}^{S} \langle v_i \rangle \ln \langle v_i \rangle \Big) \tag{S15}$$

In particular, the parameter $l$ allows to ponder the weight attributed to rare species, with $l = 1$ yielding species richness and $l = 0$ providing Shannon diversity.

The general idea is to replace relative abundances of $v_i$ by species densities $u_i$, i.e. redefining $D_l$ as

$$D_\ell = \Big( \sum_{i=1}^{S} \langle u_i \rangle^{1-\ell} \Big)^{1/\ell} \text{ for } \ell \neq 0 \text{ and } D_0 = \exp\Big( - \sum_{i=1}^{S} \langle u_i \rangle \ln \langle u_i \rangle \Big).$$

In this case, $\sum_{i=1}^{S} \langle u_i \rangle \leq 1$ and in particular $\langle u_i \rangle$ may equal zero. We thus restrict the definition of $D_l$ to $l \in [0,1]$. In addition, considering a scenario where $\langle u_i \rangle = \varepsilon / S$, we expect $D_l$ to converge to zero with ε, leading to $l \in (0,1)$. Finally, for $l = 1/2$, convergence to zero is linear in ε, and we check that $D_{1/2}$ indeed has the expected behavior in the two test scenarios. We thus consider $D_{1/2}$ throughout the simulation study.

### B. Supplementary results

##### B.1 Taking into account the dispersal of simulation outputs around their average does not alter qualitative conclusions.

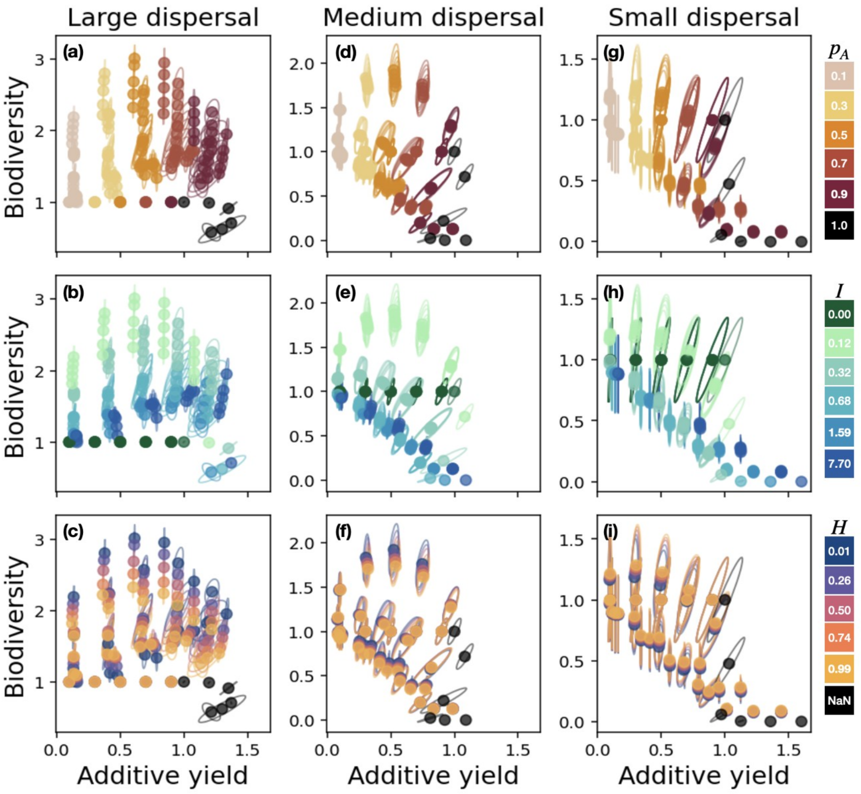


***Figure S2.*** *Average values and confidence ellipse of additive yield and biodiversity achieved by each management strategy, at large (a-c), medium (d-f) and small dispersal (g-i). Each management strategy is characterized by proportion of farmed area (a, d, g), farming intensity (b, e, h) and spatial aggregation (c, f, i). For homogeneous landscapes ($p_A$ = 1), spatial aggregation is not defined (H = NaN). For readability, the y-axis is shared only within columns.*

For each management strategy, we have performed one hundred simulations of the metacommunity dynamics. Each simulation considers one stochastic realization of the landscape with parameters *($p_A$, H, I)* characterizing the management strategy, and one stochastic sample of the species pool. For a given management strategy, biodiversity and yield are computed for each simulation, and we summarize the provided information by considering their averages (Section 2.2).

In order to take the data's scattering into consideration, we compute the confidence ellipse associated to each management strategy. The confidence ellipse is centered on the average biodiversity and yield achieved by some management strategy. Its radius equals the standard variation over all simulations for that management strategy, and its orientation is determined by the associated correlation between model outputs.

Importantly, the hierarchy obtained by ranking the management strategies according to biodiversity and additive yield is preserved (Fig. S2). Thus, discarding the information provided by the data's scattering around its mean does not alter the qualitative conclusions of Section 3.1 and 3.2.

##### B.2 Reconciliation of multiplicative yield and biodiversity is always achievable.

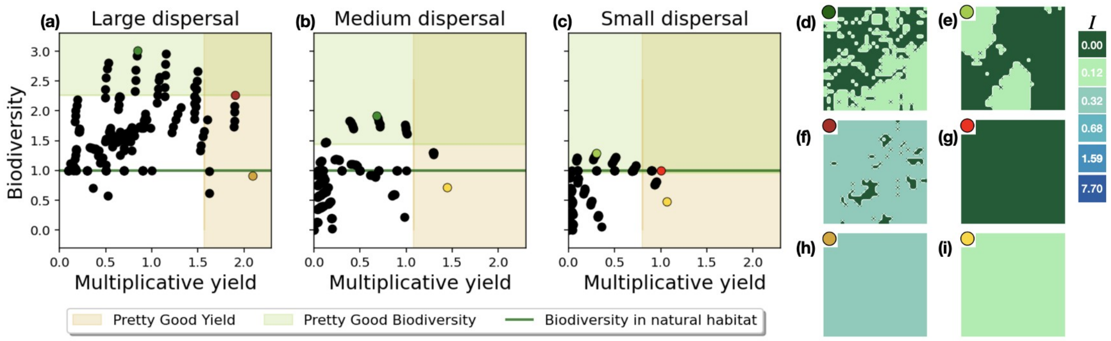


***Figure S3.*** *(A) Scatter plot of average values of multiplicative yield and biodiversity obtained for each management strategy. Orange and green shaded areas correspond respectively to areas of pretty good yield (yield superior to 75% of maximal observed yield) and pretty good biodiversity (biodiversity superior to 75% of maximal observed biodiversity). The dark green line indicates the biodiversity level achieved in a homogeneous natural environment. (B) Examples of landscapes maximizing biodiversity (d e) or multiplicative yield (h, i), as well as compromise strategies (f, g) at different dispersal levels.*

The strategies optimizing multiplicative yield or achieving a fair multiplicative yield-biodiversity compromise differ according to the species' dispersal capacity (Fig. S3). At high dispersal, farming the whole landscape at intermediate intensity maximizes multiplicative yield (Fig. S3h), and a compromise is achieved by restoring 10% of natural habitat at low spatial aggregation (Fig. S3f). At medium and low dispersal, however, multiplicative yield is best for a homogeneous low intensity farmed landscape (Fig S3i), and compromise at small dispersal requests an entirely natural habitat (Fig. S3g).

Importantly, at all dispersal levels, multiplicative yield and biodiversity collapse simultaneously as farming intensity increases (Fig. S4b, S4e, S4h). This is due to the fact that multiplicative yield strongly depends on local biodiversity (Section 2.2 and Equation S11). The level of intensity at which this breakdown occurs is lower at low dispersal capacity. At high dispersal, multiplicative yield remains strong at intermediate intensity levels (up to $I = 0.68$; Fig. S4b), whereas at medium and small dispersal, even low intensity leads to substantial multiplicative yield loss (Fig. S4e, S4h). This difference likely is due to the fact that at large dispersal, species are more robust to loss of habitat quality (Section 3.1). Thus, at large dispersal, multiplicative yield can benefit from increased farming intensity while still preserving ecosystem services. This further explains the observed differences in optimal management strategies detailed above. As for additive yield, considering the data's scattering around the mean does not change the qualitative conclusions (Fig. S5).

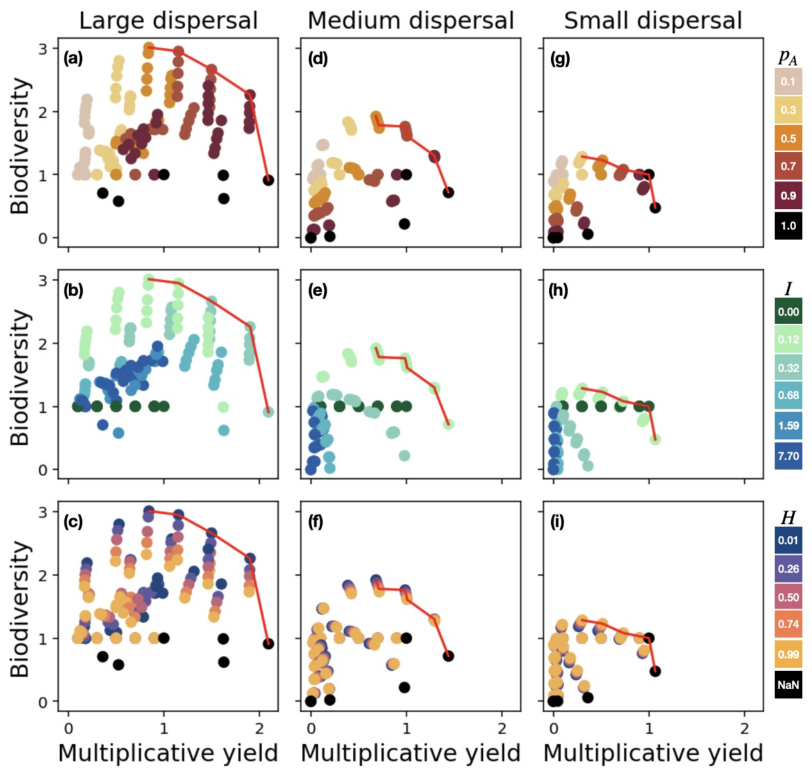


***Figure S4*** *Average values of multiplicative yield and biodiversity achieved by each management strategy, at large (a-c), medium (d-f) and small dispersal (g-i). Each management strategy is characterized by proportion of farmed area (a, d, g), farming intensity (b, e, h) and spatial aggregation (c, f, i). For homogeneous landscapes ($p_A$ = 1), spatial aggregation is not defined (H = NaN). The Pareto front is shown in red.*

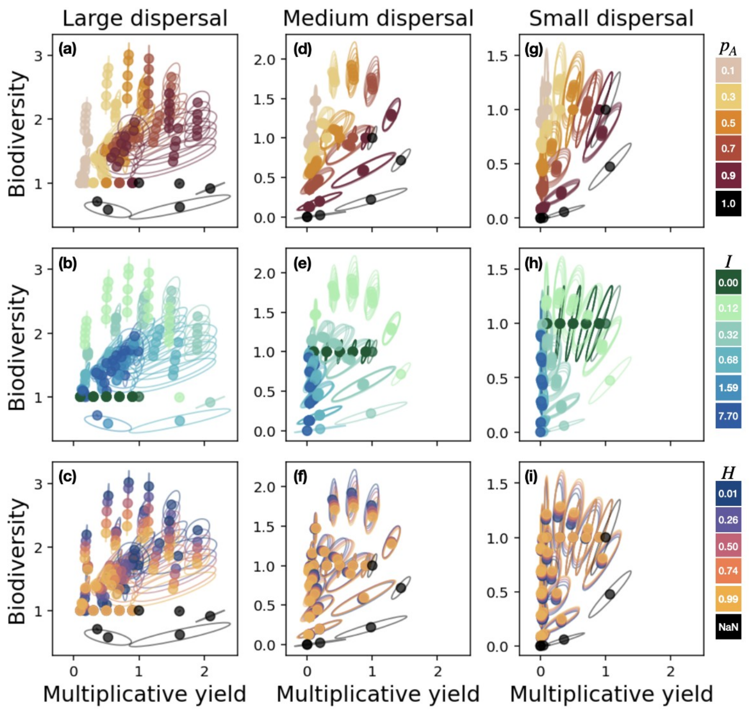


***Figure S5.*** *Average value and confidence ellipse of multiplicative yield and biodiversity achieved by each management strategy, at large (a-c), medium (d-f) and small dispersal (g-i). Each management strategy is characterized by proportion of farmed area (a, d, g), farming intensity (b, e, h) and spatial aggregation (c, f, i). For homogeneous landscapes ($p_A$ = 1), spatial aggregation is not defined (H = NaN). For readability, the y-axis is shared only within columns.*

#### Bibliography

Burian, A., Kremen, C., Wu, J.S.-T., Beckmann, M., Bulling, M., Garibaldi, L.A., *et al.* (2024). Biodiversity–production feedback effects lead to intensification traps in agricultural landscapes. *Nat Ecol Evol*, 8, 752–760.

Etherington, T.R., Holland, E.P. & O'Sullivan, D. (2015). NLMpy: a python software package for the creation of neutral landscape models within a general numerical framework. *Methods in Ecology and Evolution*, 6, 164–168.

Hill, M.O. (1973). Diversity and Evenness: A Unifying Notation and Its Consequences. *Ecology*, 54, 427–432.

Jost, L. (2006). Entropy and diversity. *Oikos*, 113, 363–375.

Jost, L. (2007). Partitioning Diversity into Independent Alpha and Beta Components. *Ecology*, 88, 2427–2439.

Loreau, M., Naeem, S., Inchausti, P., Bengtsson, J., Grime, J.P., Hector, A., *et al.* (2001). Biodiversity and Ecosystem Functioning: Current Knowledge and Future Challenges. *Science*, 294, 804–808.

Mandelbrot, B.B. & Van Ness, J.W. (1968). Fractional Brownian Motions, Fractional Noises and Applications. *SIAM Review*, 10, 422–437.

Roswell, M., Dushoff, J. & Winfree, R. (2021). A conceptual guide to measuring species diversity. *Oikos*, 130, 321–338.

Wu, D. & Xiao, Y. (2007). Geometric Properties of Fractional Brownian Sheets. *J Fourier Anal Appl*, 13, 1–37.
